# A novel real-time PCR assay panel for detection of common respiratory pathogens in a convenient, strip-tube array format

**DOI:** 10.1101/455568

**Authors:** Mohammad Rubayet Hasan, Hassan Al Mana, Virginia Young, Patrick Tang, Eva Thomas, Rusung Tan, Peter Tilley

**Author notes:** Corresponding author: Mohammad Rubayet Hasan, PhD, FCCM, D(ABMM), Clinical Molecular Microbiologist, Department of Pathology, Assistant Professor of Clinical Pathology and Laboratory Medicine, Weill Cornell Medical College in Qatar (WCMC-Q), Sidra Medicine, Office no: H2M-24093, PO Box 26999, Doha, Qatar, Direct; <u>ORCID ID: 0000-0002-4658-7949</u>.

## Abstract

Commercial multiplex assays, built on different chemistries and platforms are widely available for simultaneous detection of pathogens that cause respiratory infections. However, these tests are often difficult to implement in a resource limited setting because of high cost. In this study, we developed and validated a method for simultaneous testing of common respiratory pathogens (Respanel) by real-time PCR in a convenient, strip-tube array format. Primers and probes for sixteen PCR assays were selected from the literature or newly designed. Following optimization of individual PCR assays, strip-tube arrays were prepared by dispensing primer-probe mixes (PPM) into two sets of 8-tube strips. Nucleic acid extracts from specimens were mixed with PCR master mix, and dispensed column-wise into 2X8-wells of a 96-well plate. PPMs from strip-tubes were then added to the wells using a multichannel pipette for real-time PCR. Individual PCR assays were optimized using previously known specimens (n=397) with 91%-100% concordance with culture, DFA or PCR results. Respanel was then tested in a routine manner at two different sites using specimens (n=147) previously tested by Qiagen Resplex I&II or Fast-Track Diagnostics Respiratory Pathogens 21 assays. The sensitivity, specificity and accuracy of Respanel were 94%, 95% and 95%, respectively, against Resplex and 88%, 100% and 99%, respectively, against FTDRP21. Respanel detected 48% more pathogens (*p*<0.05) than Resplex but the rate of pathogen detection was not significantly different from FTDRP21. Respanel is a convenient and inexpensive assay that is more sensitive than Resplex and comparable to FTDRP21 for the detection of common respiratory pathogens.

## Introduction

Acute viral and bacterial respiratory tract infections are among the most common human ailments (1, 2). Symptoms range in severity from mild upper respiratory tract infections (URTI) to serious lower respiratory tract infections (LRTI), many of which are associated with significant morbidity and mortality particularly in children, the elderly and those who are immunocompromised or have underlying comorbidities such as congenital heart defect or chronic respiratory disease (3-9).

These infections contribute to an increased burden of Emergency Department visits during the winter season, leading to longer waiting times and higher health care costs, and are a common reason for hospital admission (10-13). They also account for the majority of antibiotic prescriptions, particularly in children, in spite of the fact that most of the infections will resolve on their own without medical intervention. The specific diagnosis of bacterial versus viral infection cannot conclusively be made on clinical examination alone and, as the symptoms overlap significantly, the physician may order diagnostic tests and empirically prescribe antibiotics (10).

Traditionally, the laboratory diagnosis of respiratory infections has been a lengthy procedure, using bacterial and viral culture or relatively insensitive techniques such as immunofluorescence (IFA) and rapid enzyme immune assays (EIA) (5, 11, 14-18). These test modalities can identify a limited number of organisms such as respiratory syncytial virus (RSV), influenza and parainfluenza viruses, and bacteria such as *Streptococcus pneumoniae*. With the introduction of rapid molecular tests we now know that the list of common agents known to cause respiratory infections is much longer than previously thought and the laboratory diagnostic options to detect respiratory pathogens have improved significantly (3, 6, 18, 19). Molecular tests offer many advantages including a rapid response time and the ability to detect organisms that will not grow in culture. In addition these tests allow detection of bacteria in patients where antimicrobial treatment has been initiated prior to collection of the sample. We now have access to accurate and rapid multiplex assays simultaneously detecting both viral and bacterial pathogens within a few hours. The ability to diagnose co-infections is a significant improvement as these can be associated with increased morbidity and mortality (19-21). Rapid molecular tests have been shown to reduce the length of hospital stay and the cost for testing for those with viral respiratory testing and has facilitated a more targeted approach to patients presenting with respiratory infections with respect to treatment regimens, need for admission and infection control concerns (7, 22-24).

A wide range of chemistries and platforms are now commercially available for molecular testing of common respiratory pathogens. Examples include BIOFIRE^®^ FILMARRAY^®^ Respiratory Panel (Biomerieux) that enables detection and identification of multiple organisms based on nested PCR and melt-curve analysis (25). A similar test kit, in terms of workflow and pathogen targets, is ePlex^®^ Respiratory Pathogen Panel (GenMark Dx) which makes use of signal probes and capture probes to electrochemically detect target pathogens (26). Both systems fully automate nucleic acid extraction, amplification or probe hybridization and detection. Also, these tests are very easy to perform, provide faster results (<1 hour) and require very little hands on time (~2 minutes). On the other hand, xTAG^®^ Respiratory Viral Panel (Luminex) and Resplex I&II (Qiagen) assays require nucleic acid extraction and amplification of targets prior to probe hybridization, labelling and detection by a flow cytometric method on a Luminex xMAP system (27). Multiplex, real-time PCR based kits are also available such as Fast-Track Diagnostics Respiratory pathogens 21 (FTDRP21) (Siemens Healthineers) is a five tube multiplex PCR assay to detect a total of 23 pathogen targets (28). The test kit provides reagents only and the users can run the tests using their own nucleic acid extraction platforms and real-time PCR systems.

While most commercial assays are rapid and convenient and their performance characteristics meet regulatory requirements, they are invariably expensive, which may be difficult to implement in resource poor settings. Also, commercial test kits are not uniformly available throughout the world. Timely delivery of test kits and reagents are critical for smooth operation of diagnostic laboratories and a delay in shipping may adversely affect the management of patients with respiratory infections. Furthermore, commercial tests are not free of technical limitations. Some tests may be difficult to interpret due to background signals or ambiguous results from multiple targets in the same reaction. It is also difficult to troubleshoot commercial assays because of proprietary test characteristics. In addition, commercial assays cannot be quickly modified when new pathogens or new strains of known pathogens emerge which are missed by the existing assays. Therefore, in this study, we developed an in-house PCR assay panel (Respanel) customized to detect the most common, and clinically significant respiratory pathogens. The assay was designed to use pre-aliquoted assay-specific reagents in strip-tubes for convenience, and to reduce the hands-on time required to perform multiple reactions for each patient specimen. The performance characteristics of Respanel assay were compared with two commercially available assays.

## Materials and Methods

### Clinical specimens

For individual validation of PCR assays for different pathogens, 34 reference bacterial and viral strains from American Type Culture Collection (ATCC), a total of 363 nasopharyngeal wash (NPW) and nasopharyngeal flocked swab (NPFS) specimens, 23 serum specimens, 4 pleural fluid (PF) specimens, 4 cerebrospinal fluid (CSF) specimens and 68 external quality assessment (EQA) specimens were used. These specimens were submitted to the Microbiology and Virology laboratory of BC Children’s Hospital, Canada (between January, 2012 to January, 2013) for testing either by culture or by molecular methods. For validation of the full PCR panel, 34 NPW or NPFS specimens that tested positive or negative for various pathogens by Resplex I and II (Qiagen) assays were selected from the same site. An additional 113 NPW or NPFS specimens were tested at Sidra Medicine, Qatar. These specimens include external quality assessment (EQA) specimens from College of American Pathologists (CAP), validation specimens and clinical specimens collected at Sidra Medicine from May, 2016 to August, 2017, and were previously tested for respiratory pathogens by FTD Respiratory pathogens 21 (FTDRP21) assay (Siemens Healthineers).

Resplex and FTDRP21 assays were performed according to laboratory standard operating procedures based on manufacturer’s instructions. Specimens or nucleic acid extracts were maintained at −80°C following initial testing. Testing was performed exclusively on retrospective, residual samples. To maintain patient anonymity, each sample was coded and all patient identifiers were removed to ensure that personnel involved in this study were unaware of any patient information. Ethics approval was not sought because studies that involve the secondary use of anonymous human biological materials are exempted from review by the local Research Ethics Board of the University of British Columbia and Sidra Medicine.

### Nucleic acid extraction

At BC Children’s Hospital, nucleic acids from 0.35 mL NPW or NPFS specimens were extracted using the QIAsymphony virus/bacteria kit in an automated nucleic acid extraction platform, QIAsymphony SP (Qiagen, USA). At Sidra Medicine, nucleic acids from 0.5 ml of NPW or NPFS specimens were extracted on a NucliSENS^®^ easyMAG platform (bioMérieux, France) according to the methods described by the manufacturers.

### Real-time PCR

Primers and probes for sixteen PCR assays were either obtained from previously published assays or newly designed, in this study by using the Primer Express software v3.0.1 (Life Technologies) (Table 1). The in silico specificity of the amplicon sequence for the target pathogen was confirmed by nucleotide blast search against the NCBI non-redundant nucleotide database (nr/nt) (29). Each PCR assay was individually tested and validated using reference bacterial or viral strains and clinical samples that were previously tested by a reference method such as culture, direct fluorescent antibody (DFA) tests, Resplex I & II assays (Qiagen) or an alternative PCR. Apart from Resplex assays, standard PCR assays that were available for comparison were a commercial assay for enterovirus (Trimgen Genetic Diagnostics, USA) and previously validated laboratory developed tests (LDTs) for adenovirus, influenza A, human metapneumovirus, *Streptococcus pneumoniae*, *Mycoplasma pneumoniae* and *Bordetella pertussis*,

**Table 1:**
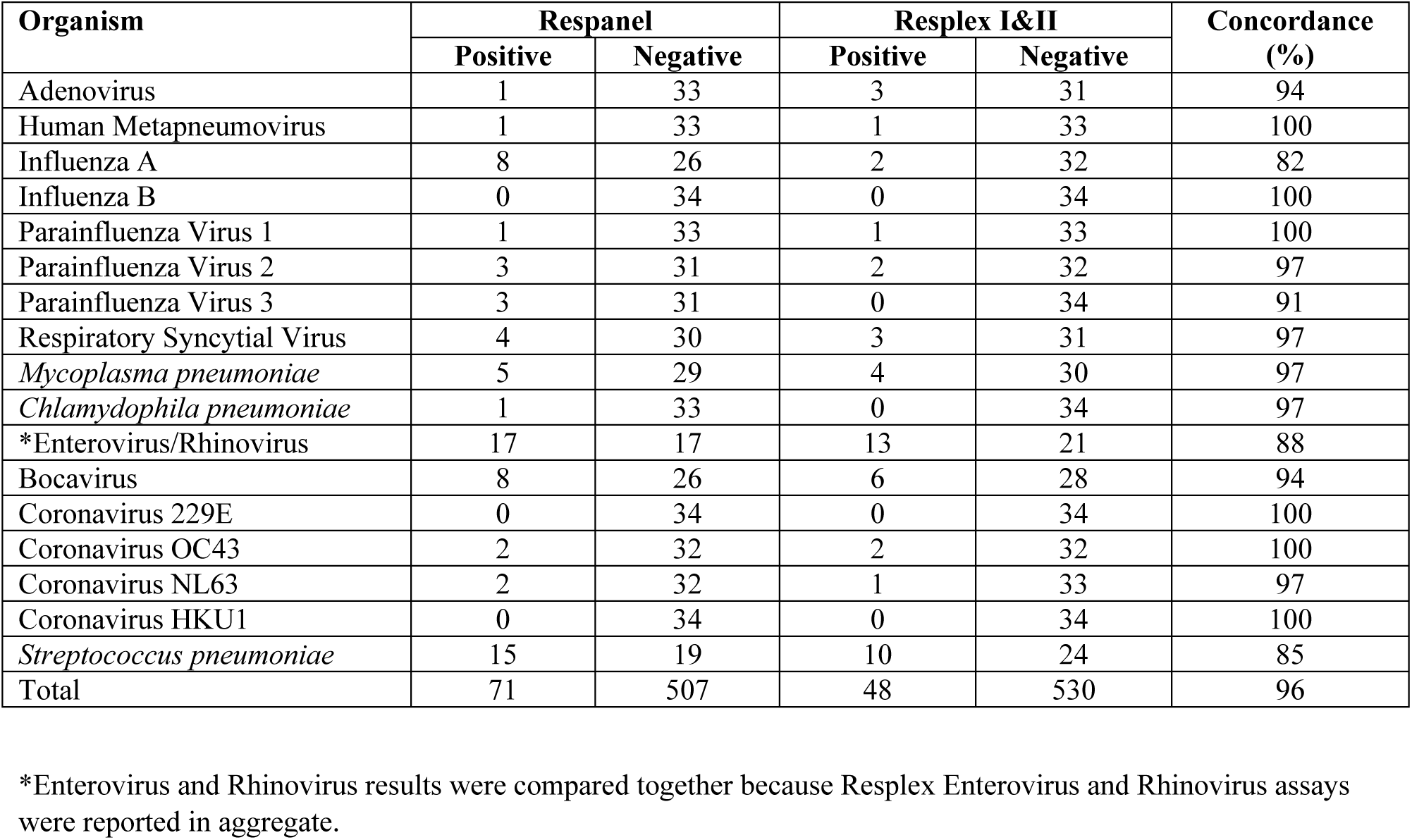
Concordance between Respanel and Resplex test results

For PCR, 5µl of sample extract was mixed with 20µl of a master mix containing 12.5 µl of 2x QuantiTect Probe RT-PCR Master Mix, 0.25 µl of QuantiTect RT Mix (Qiagen, USA) and primers and probes to final concentrations shown in Supplemental Table 1. Thermal cycling was performed in a ABI7500 Fast instrument (Thermofisher Scientific, USA) with 1 cycle of reverse transcription at 50°C for 30 min followed by 1 cycle of denaturation at 95°C for 15 min, followed by 40 amplification cycles each consisting of 94°C-15s and 60°C-60s. DNA extraction and PCR inhibition was monitored by an internal control PCR assay, using the primers and probes shown in Supplemental Table 2. For use as positive control 4 plasmids harboring amplicon sequences of all Respanel targets were custom made (Integrated DNA Technologies), diluted to 10^6^ copies/ml, mixed and aliquoted for use with each PCR runs. The aliquots of positive control were stored at −20°C. For negative controls, 0.2 ml neonatal calf serum (NCS) (Thermofisher Scientific, USA) spiked with 10^5^ copies of a plasmid, harboring target amplicon sequence of IC, were extracted along with specimens and used with each PCR runs. The target sequence of IC is a randomly generated sequence with no known homology to any sequences in the nucleotide database (29).

### Preparation of strip-tube arrays

A mixed working stock of primers and probe(s) for each of the assays were prepared to a total volume of 2 ml, according to the working stock concentrations shown in supplemental table 1. Ten µl of primer and probe mix (PPM) for each assay was dispensed into the tubes of MicroAmp Fast 8-Tube Strips (without cap) (Thermofisher Scientific, USA) according to figure 1. Strip-tubes were placed in 96-well PCR tube racks and PPMs were dispensed either with the aid of multichannel, dispensing pipettes (Eppendorf, USA) or in an automated liquid handler (Perkin Elmer, USA) into two sets of strip-tubes labeled as panel I and panel II. The strips were marked at one end with different colors for Panel I and Panel II to maintain proper orientation. Tubes were then covered with MicroAmp Optical Adhesive Film and pressed and sealed using MicroAmp Adhesive Film Applicator (Thermofisher Scientific, USA). Optical adhesive film covers were then cut vertically between the columns of tubes in each rack using a clean scalpel and panel I and II strips were stored in separate boxes at −20°C.

**Figure 1:**
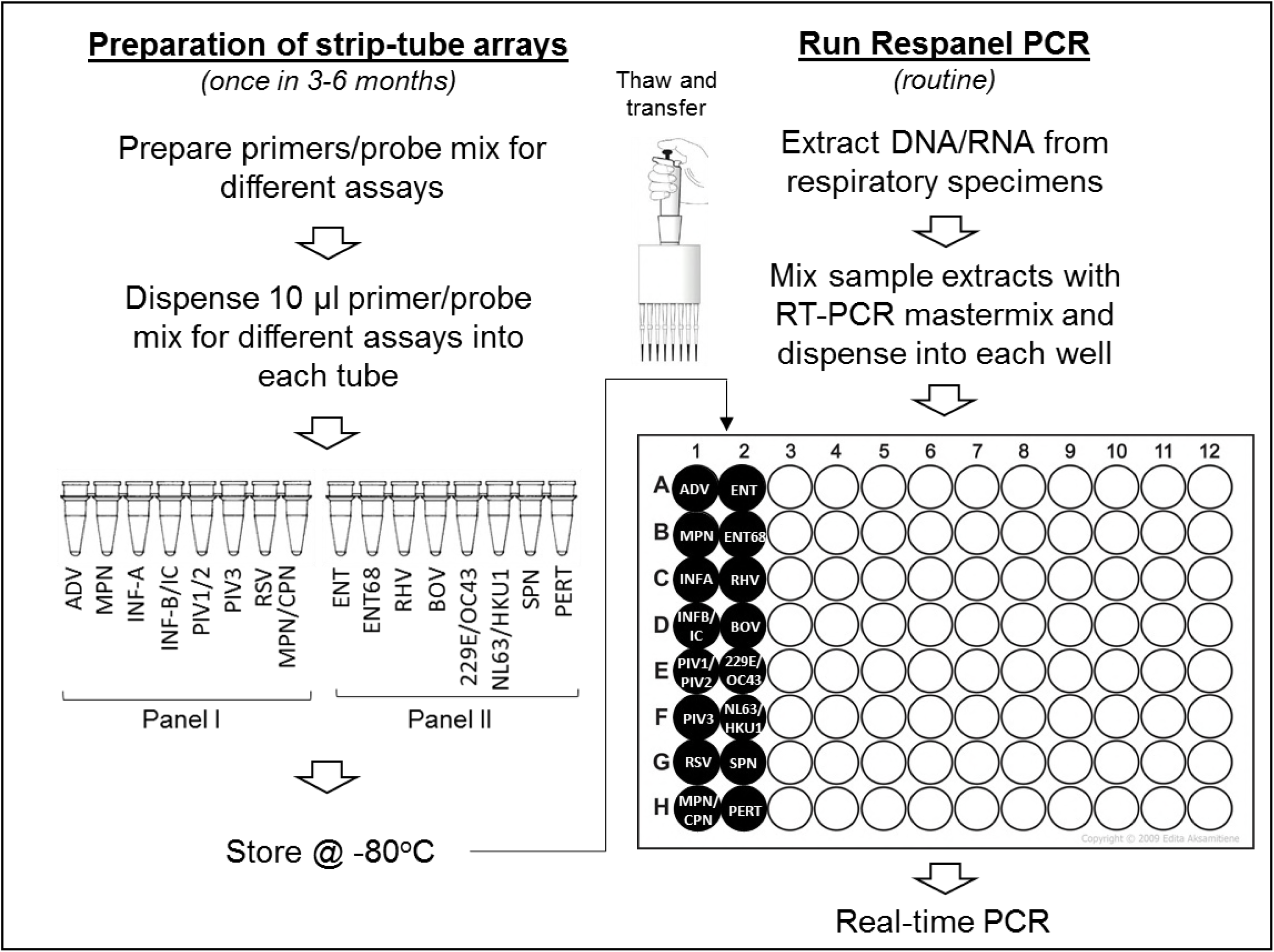
Respanel assay workflow

### Respanel PCR assay

For each sample, one panel I strip and one panel II strip are thawed, centrifuged briefly to bring the contents of the tubes at the bottom of the tubes, and placed in a 96-well PCR tube rack. A sample mix is then prepared with 88 µl of extracted nucleic acid from the specimen to be analyzed, 220 µl of 2x QuantiTect Probe RT-PCR Master Mix and 4.4 µl of QuantiTect RT Mix (Qiagen, USA). Ten percent extra volume was added to the required volume for all components, to account for pipetting error. Next, using an electronic dispensing pipette (Eppendorf, USA), 17.5 µl of sample mix was dispensed into the 16-wells of the 96-well PCR plate, in the two adjacent columns containing the primers and probes as described above and in figure 1. Alternatively, 88 µl of nucleic acid extracts were mixed with 88 µl of TaqPath™ 1-Step Multiplex Master Mix (No ROX) (Thermofisher Scientific, USA) and 44 µl of nuclease free water (Thermofisher Scientific, USA). 12.5 µl of sample mix was dispensed in the same way. 7.5 µl of each PPM from panel I and panel II strips are then transferred to the respective wells of the PCR plate using a multichannel pipette. The PCR plate was covered with adhesive film and thermal cycling was performed on an ABI7500 Fast instrument as described above (Thermofisher Scientific, USA). Test results did not vary when tests were performed in parallel with two different master mixes in a set of known specimens (data not shown).

### Statistical analysis

Correlation between the results of different assays was determined by Cohen’s kappa test. Ninety-five percent CI for sensitivity, specificity and accuracy were calculated by Clopper-Pearson interval or exact method. The significance of differences in pathogen detection rates between Respanel versus Resplex and Respanel versus FTDRP21 were calculated by 2-tailed, paired Student’s t-test.

## Results

Respanel PCR assays included singleplex assays for *Streptococcus pneumoniae*, *Bordetella pertussis*, adenovirus, human metapneumovirus, influenza A, parainfluenza virus 3, respiratory syncytial virus, enterovirus, rhinovirus, and bocavirus and duplex assays for *Mycoplasma pneumoniae* and *Chlamydia pneumoniae*, influenza B and an internal control (IC), parainfluenza virus 1 and parainfluenza virus 2, coronavirus 229E and coronavirus OC43, and coronavirus NL63 and coronavirus HKU1, respectively. Test comparisons were only made between results for matching pathogens. Resplex assays included all Respanel targets except *Bordetella pertussis* and also incuded *Neisseria meningitidis*, *Haemophilus influenzae* and *Legionella pneumophila* that are not included in Respanel. Similarly, FTDRP21 assays include influenza A(H1N1)pdm09 virus, human parainfluenza virus 4, human parechovirus, *Haemophilus influenzae* B and *Staphylococcus aureus* that are not included in Respanel.

PCR conditions for all targets included in Respanel were first optimized using ATCC strains or EQA panels. Primer and probe concentrations were adjusted for maximum analytical sensitivity and specificity (Supplemental table 2). In some cases such as Influenza B and enterovirus, multiple primer and probe sets were tested for superior performance. Because of the cross-reactivity of enterovirus and rhinovirus primers and probes with their respective targets, an additional PCR reaction was added for enterovirus 68 in order to correctly assign results for enteroviruses and rhinoviruses. PCR assays were then tested individually using PCR, DFA or culture confirmed specimens or ATCC reference strains of bacteria and viruses (n=397). Respanel PCR results in 91%-100% agreement with the reference results (Supplemental table 3).

Next, a total of 34 nasopharyngeal wash specimens that were previously tested by Resplex I and II were tested by Respanel assay as described in the materials and methods. The overall, observed agreement with Resplex assay was ~95% (*kappa* = 0.734; 95%CI: 0.643-0.826). In total, 71 pathogens were identified in these samples by Respanel, which was 48% higher than that of Resplex assays (Table 1). >90% of pathogens that were undetectable by the Resplex assays had higher C_T_ values (>30) by real-time PCR. Similarly, 113 NPW or NPFS specimens that were previously tested by FTDRP21 assay were retested by Respanel. The overall, observed agreement with FTDRP21 assay was ~99.2% (*kappa* = 0.896; 95%CI: 0.845-0.946). In total, 78 pathogens were identified in these samples by Respanel and 82 pathogens were identified by FTDRP21 assay (Table 2). Respanel detected 6 pathogens that were not detected by FTDRP21. On the other hand FTDRP21 assay detected 10 pathogens that were not detected by Respanel. Notably, majority of these specimens had PCR C_T_ values >35.

**Table 2:** Concordance between Respanel and FTD Respiratory pathogens 21 test results

| Organism | Respanel |  | FTD |  | Concordance (%) |
| --- | --- | --- | --- | --- | --- |
|  | Positive | Negative | Positive | Negative |  |
| Adenovirus | 7 | 106 | 7 | 106 | 100 |
| Human Metapneumovirus | 2 | 111 | 3 | 110 | 99 |
| Influenza A | 4 | 109 | 5 | 108 | 99 |
| Influenza B | 3 | 110 | 3 | 110 | 100 |
| Parainfluenza Virus 1 | 3 | 110 | 3 | 110 | 100 |
| Parainfluenza Virus 2 | 2 | 111 | 2 | 111 | 100 |
| Parainfluenza Virus 3 | 5 | 108 | 6 | 107 | 99 |
| Respiratory Syncytial Virus | 8 | 105 | 8 | 105 | 100 |
| *Mycoplasma pneumoniae* | 4 | 109 | 4 | 109 | 98 |
| *Chlamydomphila pneumoniae* | 0 | 113 | 0 | 113 | 100 |
| Enterovirus | 4 | 109 | 4 | 109 | 98 |
| Rhinovirus | 19 | 94 | 21 | 92 | 96 |
| Bocavirus | 4 | 109 | 4 | 109 | 98 |
| Coronavirus 229E | 3 | 110 | 4 | 109 | 99 |
| Coronavirus OC43 | 3 | 110 | 3 | 110 | 100 |
| Coronavirus NL63 | 5 | 108 | 4 | 109 | 99 |
| Coronavirus HKU1 | 2 | 111 | 1 | 112 | 99 |
| Total | 78 | 1843 | 82 | 1839 | 99 |

The sensivitivity and specificity of Respanel assay and the accuracy of results obtained by this assay were calculated against both Resplex and FTDRP21 assays. The specificity and accuracy of Respanel results were ≥95% against both commercial assays. Sensitivity of Respanel was 94% and 88% against Resplex and FTDRP21assays, respectively (Table 3). When results for each of the individual pathogens were compared, Respanel assay results were more correlated (>95%) to FTDRP21 assay than Resplex assays (Table 1 and 2).The cost of Respanel is approximately 25% of Resplex assays and requires less than 10 minutes of hands-on time per sample, apart from the initial effort required for producing PCR-ready aliquots of primer/probe mixes in PCR strip-tubes.

**Table 3:**
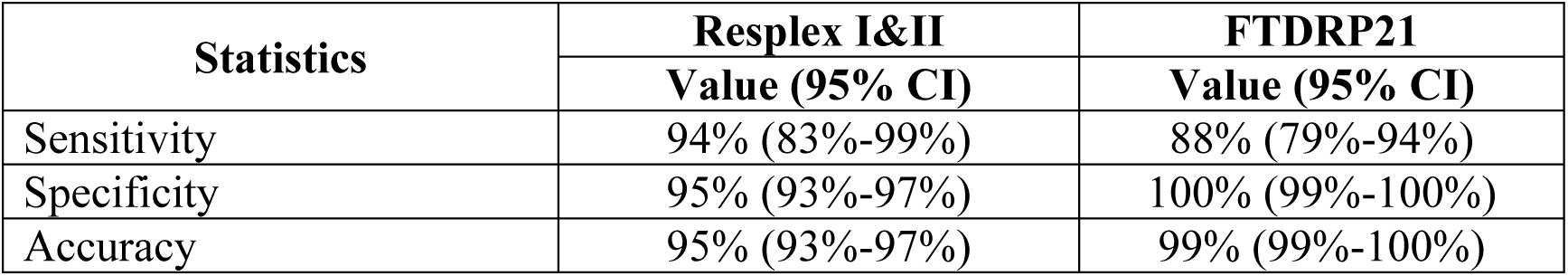
Diagnostic performance of Respanel compared to Resplex and FTD Respiratory pathogens 21 assays

| Statistics | Resplex I&II | FTDRP21 |
| --- | --- | --- |
|  | Value (95% CI) | Value (95% CI) |
| Sensitivity | 94% (83%-99%) | 88% (79%-94%) |
| Specificity | 95% (93%-97%) | 100% (99%-100%) |
| Accuracy | 95% (93%-97%) | 99% (99%-100%) |

## Discussion

Molecular assays for simultaneous detection of common respiratory pathogens are now widely used, replacing many conventional diagnostic methods such as culture and antigen detection assays. Implementation of these tests have resulted in increased identification of causative pathogens for acute upper and lower respiratory tract infections. In addition, a significant decrease in TAT has assisted institutions in the timely use appropriate isolation precaution measures and reduce inappropriate antibiotic use (30-33). A wide range of multiplex assays are now available commercially which differ in their use of chemistry, optics and the level of automation. Many test methods have also been described in the literature for potential use as laboratory-developed tests (LDT). To make an appropriate choice, a careful assessment of performance characteristics as well as costs and benefits associated with the test method is necessary. Ideally, when performance characteristics of test methods are comparable and meet regulatory requirements, a test method and a platform that provides rapid TAT and requires minimum technical skill and hands on time would be preferable. However, the costs associated with the platform, such as test kits and service contracts may not be affordable to many laboratories. Furthermore, because the majority of these test platforms are closed systems it is difficult to troubleshoot ambiguous results and test failures in a timely manner without active and continuous support from the vendor. LDTs on the other hand necessitate more labour and responsibilities for the laboratory in order to maintain the quality of the test. Substantial molecular biology expertise and effort are also necessary for validation of LDTs. However, LDTs are less expensive and can be tailored specifically to needs of the laboratory and the patient population it serves. An additional advantage is that LDTs can be updated rapidly to make the test inclusive for a new pathogen of interest or an emerging variant of a known pathogen. Logistically, if stored properly, primers and probes used in PCR based LDTs may have a longer shelf life than the manufacturer recommended shelf life of commercial kits.

The purpose of this study was to develop a laboratory-developed, real-time PCR based test panel for selected respiratory pathogens with minimum multiplexing and in a format so that minimum hands-on time is required. Accordingly, we designed our workflow so that assay specific reagents are ready to be used for routine testing (Figure 1). We also minimized repeated pipetting through the use of multichannel pipettes for faster transfer of reagents and to prevent pipetting error. The pre-aliquoted primer and probe mixes in 8-tube strips can be prepared once in 3 months, 6 months or 1 year according to laboratory needs. Preparation of primer/probe mixes and strip tubes for Panel I and II may take one FTE technologist time for up to 8 hours. We noted that in an automated liquid handler (Perkin Elmer, USA), preparation of 200 test strips takes about 2 hour time.

After initial optimization of PCR conditions, Respanel assay was validated in three steps. First, the accuracy of results obtained by each PCR assay was compared with results obtained by bacterial or viral culture, DFA, Resplex and standard PCR assays offered by the Microbiology and Virology laboratories of BC Children’s Hospital, Canada. Next test comparisons were made with Resplex and FTDRP21 assays at two different laboratories using nasopharyngeal specimens. Despite multiple test types used as reference methods, Respanel assay demonstrated 85%-100% concordance with the reference methods. Respanel detected significantly (p=0.015 by Student’s T-test) more pathogens than Resplex but was not different than the FTDRP21 panel (p=0.27 by Student’s T-test). This is consistent with the fact that Respanel and FTDRP21 are both real-time PCR based methods, while Resplex assay employed a different chemistry based on highly multiplexed PCR, probe hybridization and flow cytometry.

To our knowledge, this is the first report of a strip-tube array based, clinical laboratory developed PCR test panel for respiratory pathogens designed for cost savings, convenience and superior or equivalent performance compared to commercial assays. Furthermore, the laboratory developed, Respanel has the additional advantage of flexibility to update, modify, incorporate or replace pathogen targets as required, and to serve specific patient populations with acute respiratory infections.

## Acknowledgements

This research received no specific grant from any funding agency in the public, commercial, or not-for-profit sectors.

